# Genome sequence and characterisation of coliphage vB_Eco_SLUR29

**DOI:** 10.1101/787416

**Authors:** Ibrahim Besler, Pavelas Sazinas, Christian Harrison, Lucy Gannon, Tamsin Redgwell, Slawomir Michniewski, Steven P. Hooton, Jon L. Hobman, Andrew Millard

## Abstract

Bacteriophage that infect *Escherichia coli* are relatively easily isolated, with greater than 600 coliphage genomes sequenced to date. Despite this there is still much to be discovered about the diversity of coliphage genomes. Within this study we isolated a coliphage from cattle slurry collected from a farm in rural England. Transmission electron microscopy identified the phage as member of the *Siphoviridae* family. Phylogenetic analysis and comparative genomics further placed it within the subfamily *Tunavirinae* and forms part of a new genus. Characterisation of the lytic properties reveals that it is rapidly able to lyse its host when infected at high multiplicity of infection, but not at low multiplicity of infection.

## Introduction

Bacteriophages infecting *Escherichia coli* (coliphages) are readily isolated from a variety of sources, with > 600 complete or near complete coliphage genomes publicly available (May 2019). Coliphage genomes range in size from 3.39 kb ^1^ to 386.44 kb ^2^ and are represented in a number of phage families including the *Leviviridae*, *Siphoviridae*, *Podoviridae*, *Myoviridae* and the *Microviridae*. Currently within these families, there are 37 genera and 157 species that contain coliphages, with the many taxa being poorly sampled ^3^. Therefore, there is still much to be discovered by the continued isolation and sequencing of coliphage, with many new phage types and taxa still to be discovered ^3^. Building on our previous research that has isolated coliphages from animal slurries ^4,5^, we aimed to further isolate phage from this system and characterise the phenotypic properties.

## Materials and Methods

Bacteriophage vB_Eco_SLUR29 was isolated from cattle slurry that was collected from a farm in the East Midlands, in the United Kingdom. The double-agar overlay method was used for phage isolation and the subsequent three rounds of purification using *E. coli* K-12 MG1655 as host, as has been described previously. High titre lysate was produced by infection of ~50 ml of exponentially growing *E. coli* MG1655 and incubated at 37°C with shaking at 300 rpm, until lysis had occurred. Phage growth parameters (burst size, eclipse and latent period) were determined by performing one-step growth experiments as described by Hyman and Abedon (2009), with free phages being removed from the culture by pelleting the host cells via centrifugation at 10,000 g for 1 min, removing the supernatant and re-suspending cells in fresh medium ^6^. Three independent replicates were carried out for each experiment. The virulence index was calculated using the method described by Storms and Sauvageau (2019) using a SPECTROstar Omega (BMG) plate reader. For genome sequencing DNA was extracted from 1 ml of bacteriophage lysate as previously described ^7^. One nanogram of DNA was uses as input for Nextera XT library preparation, following the manufacturer’s instructions. Sequencing was performed on an Illumina MiSeq (250 bp paired-end) with both the genome sequence (accession: LR596614) and raw reads (accession: ERR3385641) submitted to the ENA under Project accession: PRJEB32519

### Bioinformatics and comparative genomics

Reads from sequencing were trimmed with Sickle v1.33 using default parameters ^8^. Assembly was carried out with SPAdes v.3.6.0 using “--only-assembler” option ^9^. Genome assembly errors were corrected with two rounds of checking and polishing with Pilon v1.23 using default parameters ^10^. The genome was annotated with Prokka 1.12 using a protein database constructed from accession LR027385 ^11^. The start of the genome was arbitrarily set at the gene encoding for the large terminase subunit, for ease of comparison. For rapid comparison against all other phage genomes a Mash database was constructed of all complete bacteriophage genomes available at the time of analysis (~ 11,000, April 2019) using the following Mash setting: “ –s 10000” using a previously described method ^12^. Closely related genomes were identified using this database and the `dist` function. From this initial set of genomes, the *terL* gene was used to construct a phylogenetic tree using IQ-TREE^13^. Following this, a more detailed analysis of the most closely related genomes was carried out. Phage genomes that were found to be similar were re-annotated with Prokka to ensure consistent gene calling between genomes for comparative analysis ^11^ and the GET_HOMOLOGUES pipeline used to identify core genes ^14^. For calculation of phage average nucleotide identity, pyani was used with default settings ^15^. Core gene analysis for phages within the putative genus *Swanvirus* was carried out with ROARY using “--e --mafft -p 32 –i 90” ^16^.

### Transmission Electron Microscopy

TEM was carried out at the University of Leicester Core Biotechnology Services Electron Microscopy using a previously described method ^12^. Digital images were collected with a Megaview III digital camera using iTEM software. Phage images were processed in ImageJ using the measure tool, with the scale bar present used as a calibration to measure phage particle size ^17^. The data presented in the mean of 15 phage particles.

## Results

Bacteriophage vB_Eco_SLUR29 was isolated from animal slurry collected from a farm in the East Midlands, United Kingdom using *E. coli* K-12 MG1655 as a host. The plaque morphology was small (< 3 mm) clear plaques, suggestive of an obligately lytic phage. Imaging of phage vB_Eco_SLUR29, using transmission electron microscopy, revealed a polyhedral head with a long flexible non-contractile tail. The head was 57 nm (± 2.8) and 56.7 nm (± 6.7) in width and length respectively, with a tail that was 12 nm (± 1.4) wide and 142 nm (± 23.2) long. The long non-contractile tail allowed its classification within the *Siphoviridae* and the head length:width ratio further classified vB_Eco_SLUR29 within subgroup B1 ^18^ (Fig. 1A).

**Figure 1.**
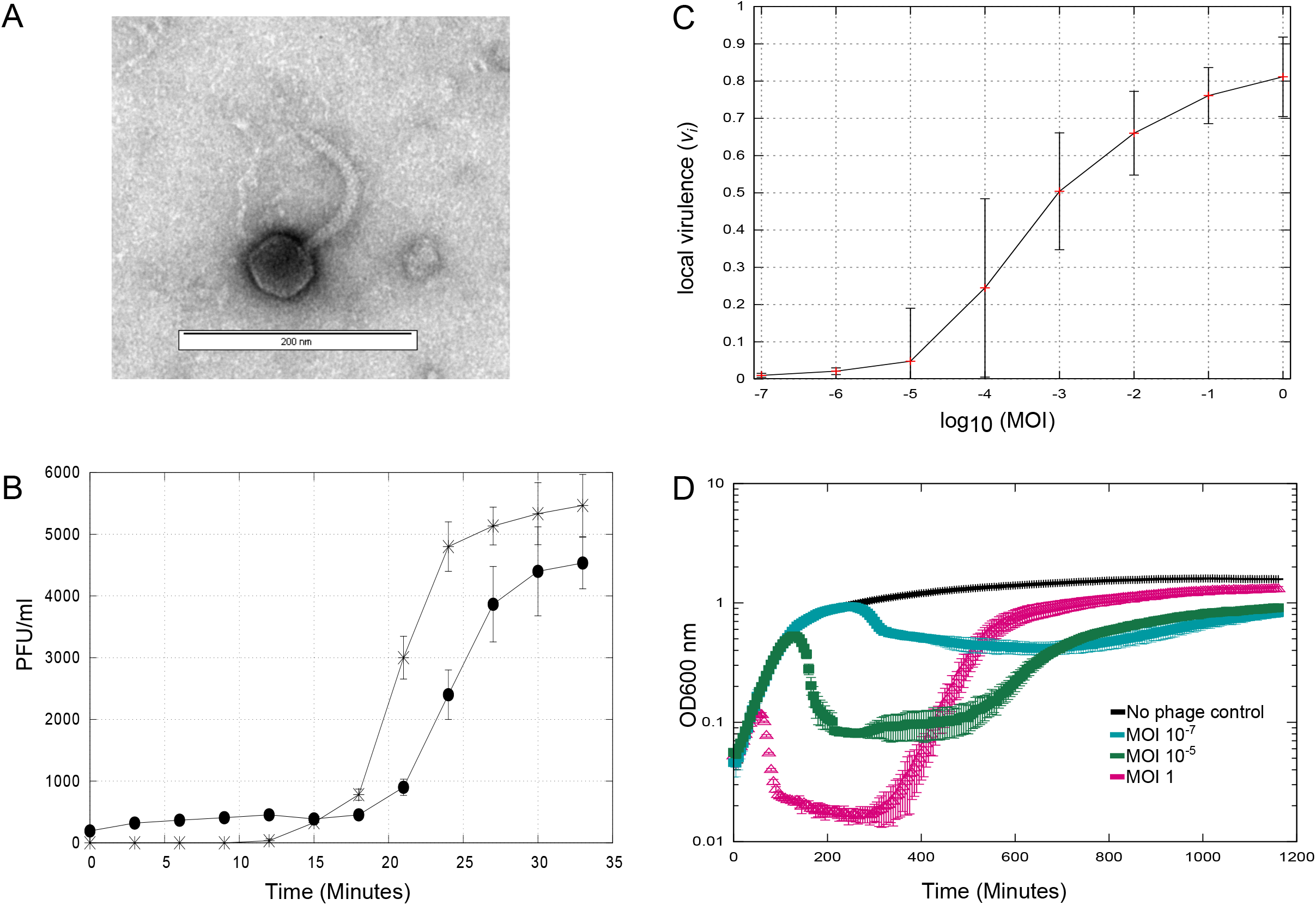
Phenotypic properties of phage vB_Eco_SLUR29. A) TEM image of phage vB_Eco_SLUR29, identifying the morphological features representative of a siphoviruses. B). One-step growth experiment (n=3), samples treated with CHCl_3_ are closed black circles, untreated samples are stars. C) Local virulence index of vB_Ecol_SLUR29 at MOIs ranging from 1 - 1×10^−7^. D) Killing curves of vB_Eco_SLUR29 against *E .coli* K-12 MG1655 (n=3).

The lytic properties of the phage vB_Eco_SLUR29 were determined using a one-step experiment (Fig 1B). The latent period was determined to be ~ 21 mins, the eclipse period 17 mins, with a burst size of 25 (+/-6). To further characterise the infection properties, killing curves were used to investigate the ability of vB_Eco_SLUR29 to kill its host over a range of MOIs and also determine the recently described virulence index ^19^. At an MOI of 1, there was rapid lysis of the culture with near complete lysis after 100 minutes, followed by a steady increase in growth after 300 minutes. When a very low MOI of 1×10^−5^or lower was used, these cultures displayed growth comparable to an un-infected control until the onset of lysis was observed at ~300 minutes (MOI 1×10^−7^). Post 300 minutes a decrease in growth was observed, before a steady increase from 700 minutes onwards. For calculation of the virulence index, the local virulence values at MOIs from 1 to 10^−7^ were determined (Fig 1C). This further highlighted that phage vB_Eco_SLUR29 was inefficient at lysing its host at very low MOIs (Fig 1C). The virulence index of phage vB_Eco_SLUR29 was calculated to be 0.37 at 37 ^°^C in LB medium.

### Genome Sequencing

To further classify vB_Eco_SLUR29 beyond morphological similarity to other Siphoviruses, the genome of vB_Eco_SLUR29 was sequenced. Genome sequencing resulted in a single chromosome with an average coverage of 426x. The genome was 48,466 bp in length with a G+C content of 44.7 %, with 78 predicted genes and no tRNAs. Of the 78 predicted genes, functions could only be predicted for 25 of the proteins they encode and the majority of these were phage structural proteins. Comparing the genome sequence of vB_Eco_SLUR29 against current phage genomes identified that it had the greatest similarity (Mash distance < 0.05) to the coliphages SECphi27, vB_Eco_swan01, vB_EcoS-95, vB_Eco_mar001J1 and vB_Eco_mar002J2. These phages have previously been found to form a monophyletic cluster, which represents a putative genus within the subfamily Tunavirinae ^12^. In addition, vB_Eco_SLUR29 showed some similarity to a group of phages infecting *Campylobacter* (Mash distance < 0.25), that are not currently classified by the ICTV.

A phylogeny was reconstructed using *terL*, with the top 100 hits to the vB_Eco_SLUR29 *terL* based on BLASTP analysis. The resultant phylogeny placed vB_Eco_SLUR29 within the subfamily Tunavirinae, forming a clade with the coliphages vB_Eco_swan01 (acc: LT841304)^20^, SECphi27 (acc: LT961732), vB_Eco_mar001J1 (acc: LR027388), vB_Eco_mar002J2 (acc: LR027385) and vB_EcoS-95 (acc: MF564201)^21^ which is a sister group to the clade containing phage pSF-1, the sole representative of the genus *Hanrivervirus* ^22^. To further clarify the position of vB_Eco_SLUR29 within the subfamily Tunavirinae, a core-gene phylogeny was constructed for phages that were closest to vB_Eco_SLUR29. This included phages of the genera *Tlsvirus, Webervirus*, *Hanrivervirus* and the large group of unclassified phages that infect *Campylobacter* (Fig 2). Four core genes were identified in this set of phage (representative homologues are: SLUR29_0019, SLUR29_0039, SLUR29_0053 and SLUR29_0059) using the GET_HOMOLOGUES pipeline^14^. The resultant four genes were concatenated and used in further phylogenetic analysis. This phylogeny confirmed that phage vB_Eco_SLUR29 formed a clade with the coliphages vB_Eco_swan01 SECphi27, vB_EcoS-95, vB_Eco_mar001J1 and vB_Eco_mar002J2 all of which were isolated on *E.coli* and likely represent a new genus. The *Shigella* infecting phage pSf-1 again grouped with phage A16a, with this cluster being more closely related to the group of unclassified phages infecting Campylobacter than it is to vB_Eco_SLUR29.

**Figure 2.**
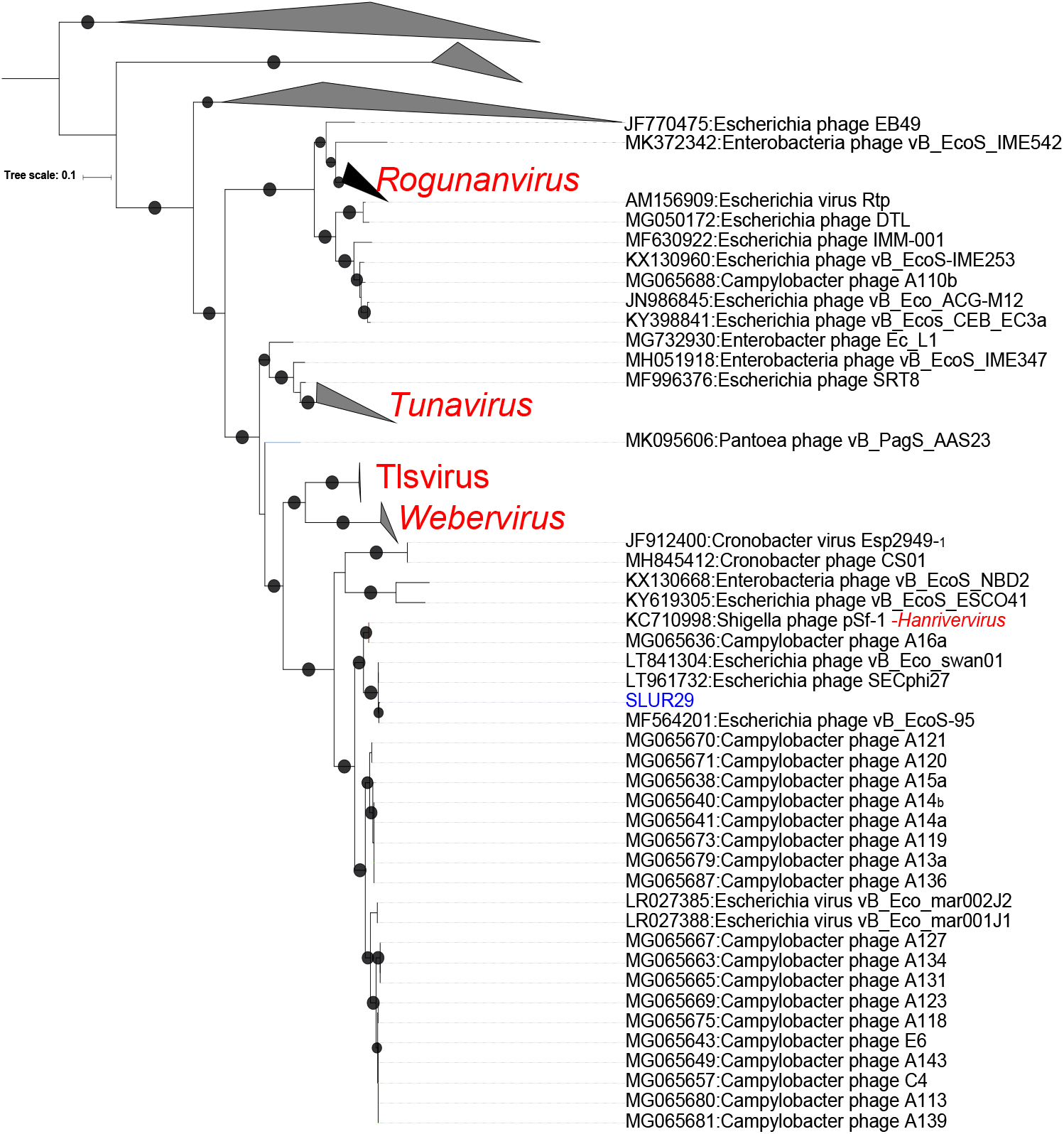
Phylogenetic analysis of vB_Eco_SLUR29. The phylogeny was created using the *terL* gene as marker. Sequences were aligned with MAFTT ^24^ and trees constructed with IQ-TREE with 1000 bootstrap replicates^13^. Bootstrap values > 70 are marked with a black circle, with increasing size proportional to the bootstrap values. The clades containing phages of the genera *Tlsvirus* and *Webervirus* were collapsed for clarity.

### Comparative Genomics

Phages vB_Eco_SLUR29, vB_Eco_swan01, SECphi27, vB_EcoS-95, vB_Eco_mar001J1, vB_Eco_mar002J2 all have average nucleotide identity (ANI) above 90% with each other and an ANI of < 80% with pSF1 (Fig 3). Whole genome comparisons of vB_Eco_SLUR29, vB_Eco_swan01, SECphi27, vB_EcoS-95, vB_Eco_mar001J1, vB_Eco_mar002J2 reveals they have 51core genes (Fig 4) and have high degree of synteny across the genomes (Fig 4). Inclusion of the phage pSF-1 in this analysis, results in only 1 core-gene.

**Figure 3.**
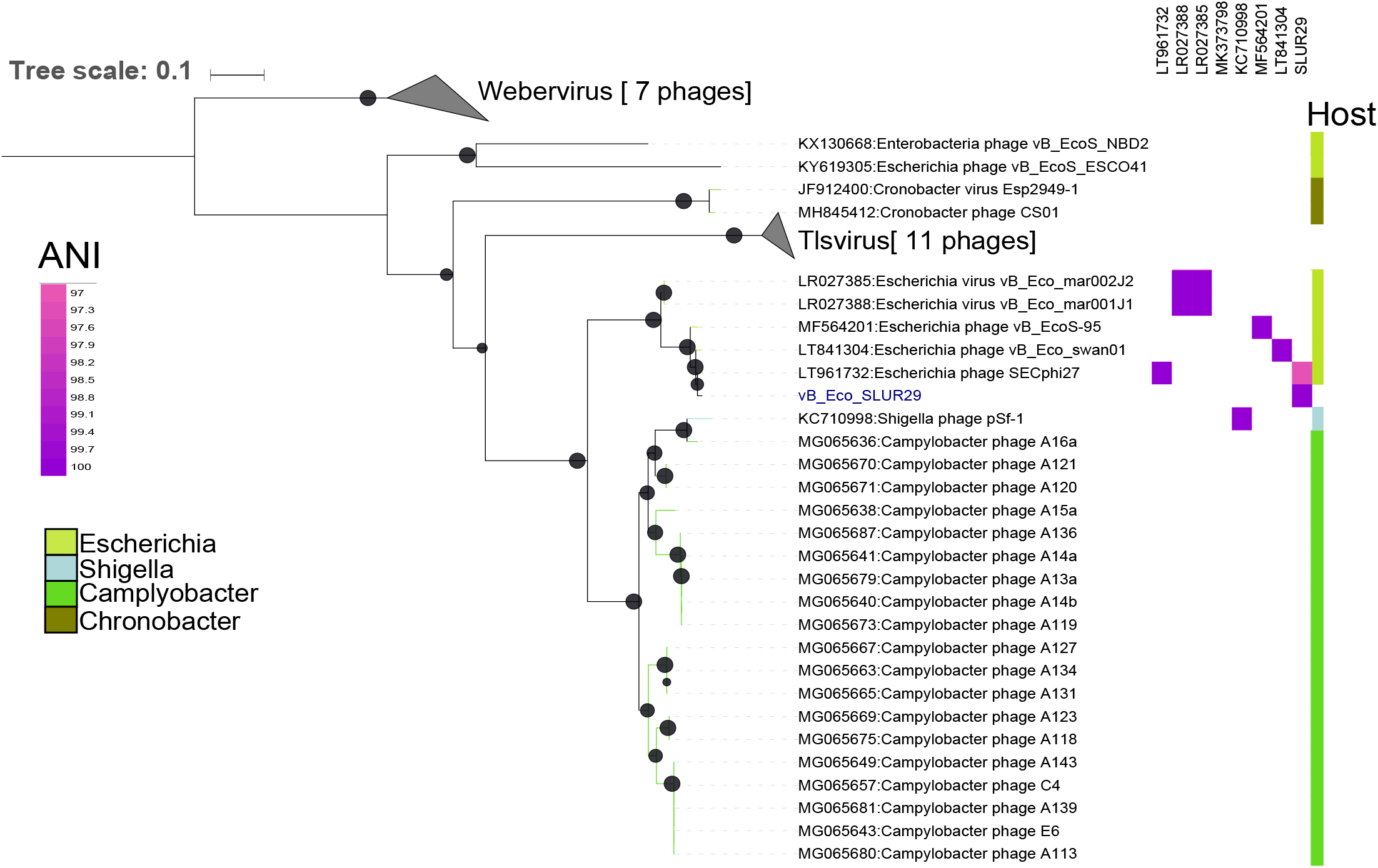
Phylogenetic analysis of vB_Eco_SLUR29. The phylogeny was created using four concatenated core-genes. Genes were aligned with MAFFT ^24^ and tress constructed with IQ-TREE ^13^, using a SYM+R4 model of evolution. Bootstrap values > 70 are marked with a black circle, with increasing size proportional to the boostrap values. The clades containing phages of the genera *Tlsvirus* and *Webervirus* were collapsed for clarity in presenting the tree.

**Figure 4.**
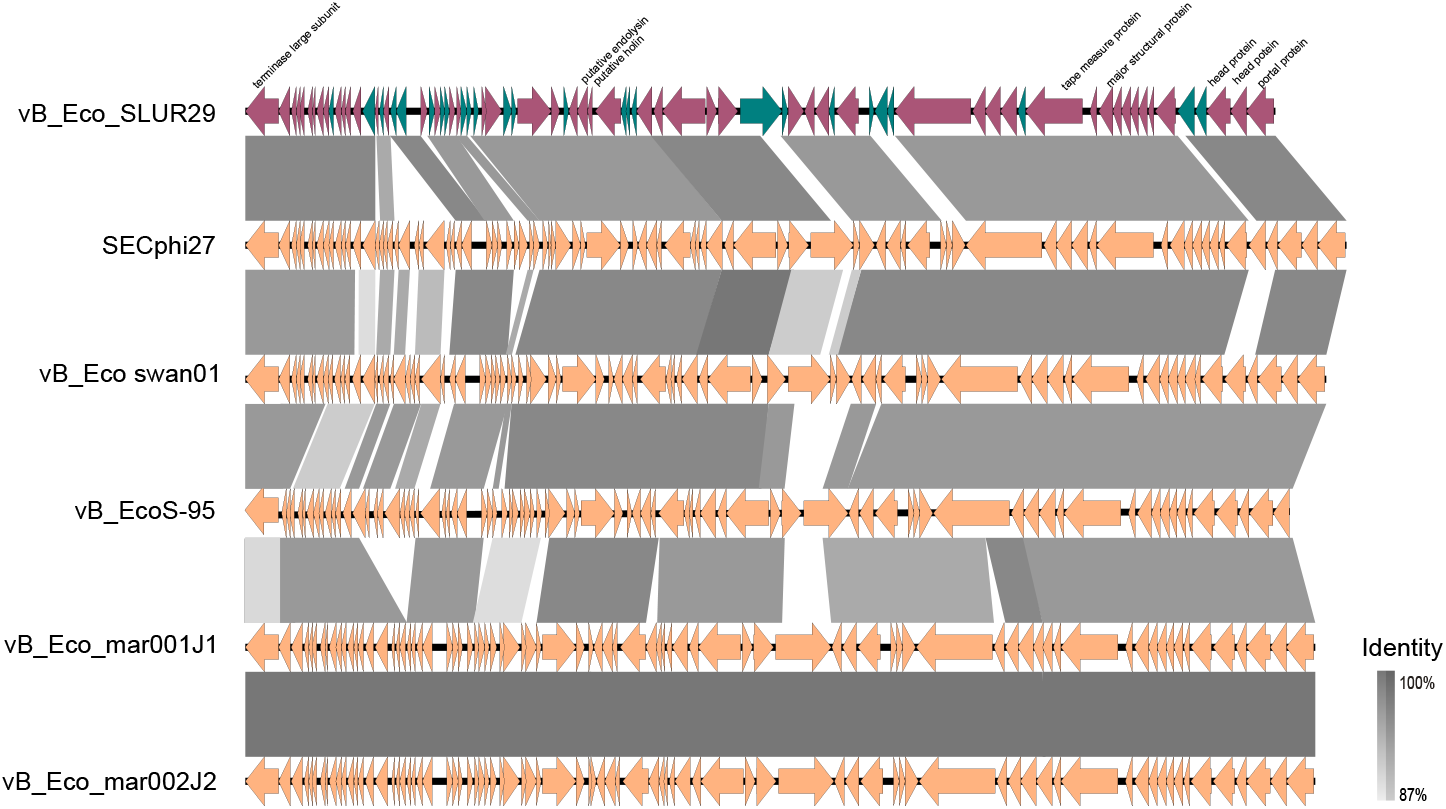
Comparative genomics of vB_Eco_SLUR29 with phages SECphi27, vB_Eco_swan01, vB_EcoS-95, vB_Eco_mar001J1 and vB_Eco_mar002J2. Alignments were constructed with EasyFig using blastn ^25^. Selected genes are annotated for vB_Eco_SLUR29. Genes that are coloured purple and green are core-genes and accessory respectively, and are only marked on phage vB_Eco_SLUR29.

## Discussion

We have previously isolated coliphages from the same slurry tank and have found phages that are representatives of the genera *T4virus* and *Seuratvirus* ^4,5^. This is the first report of phage from within the subfamily *Tunavirinae* from this particular slurry tank environment. Wether this is a reflection of their abundance or due to small sample sizes remains to be seen. Comparison of vB_Eco_SLUR29 with its closest relatives, reveals they have all been isolated on *E.coli* K-12 MG1655, although many can infect other bacteria within the Enterobacteriaceae (Fig 3)^12,21^.

Given the high sequence identity between vB_Eco_SLUR29 and its closest relatives, it was unsurprising that it has similar morphological properties when examined by transmission electron microscopy. Phylogenetic analysis clearly places vB_Eco_SLUR29 within the subfamily *Tunavirinae*, in a clade that is sister to a clade that contains phages of the genus *TLsvirus* and contains the phages vB_Eco_mar001J1, vB_Eco_mar002J2, vB_EcoS-95, vB_Eco_swan01 and SECphi27. Previously, we have suggested SECphi27, vB_Eco_mar001J1, vB_Eco_mar002J2, vB_Eco_swan01 and pSf1 are members of the same genus ^12^. Since then *Shigella* phage pSf1 has been formally classified as the sole member of the genus *Hanrivervirus.* The addition of phages vB_Eco_SLUR29, vB_EcoS-95 and *Campylobacter* infecting phages to the database further clarifies the position of *Shigella* phage pSf1, into a separate clade from vB_Eco_mar001J1, vB_Eco_mar002J2 and SECphi27. The current starting point for classification of phage species is >95 % ANI^23^. Using this criterion, vB_Eco_SLUR29, vB_Eco_mar001J1, vB_Eco_mar002J2, vB_EcoS-95, vB_Eco_swan01, SECphi27 and vB_EcoS-95 would form a single species. This is inconsistent with the phylogeny observed and a higher ANI cut-off of >97% might be more suitable for this group of phages, as has previously been suggested for this and other phage groups ^5,12^. Thus we propose this group of phages represents a new genus with four species, represented by the type phages vB_Eco_mar001J1, vB_EcoS-95, vB_Eco_swan01 and SECphi27. With vB_Eco_SLUR29 being the same species as phage SECphi27, which was isolated first. We propose the genus is named *Swanvirus* after the phage vB_Eco_swan01, which was the first isolated phage in the genus.

All phages of the proposed genus *Swanvirus*, have a conserved and syntenous genome structure. In contrast to the conservation of genes between phages, there is greater variation in phenotypic properties. The burst size of vB_Eco_SLUR29 is 22, which is smaller than the reported values of 115, 78 and 51 for vB_Eco-S59, vB_Eco_swan01 and vB_Eco_mar002J2 respectively ^12,21^. There is also considerable variation in the latent period for these phages, with vB_Eco_SLUR29 having a substantially longer latent period (21 min) than the reported 4, 12 and 15 min for vB_Eco-S59, vB_Eco_swan01 and vB_Eco_mar002J2 ^12,21^. However, the reported 4 min latent period of vB_Eco-S59 seems extraordinarily rapid and maybe an artefact of the method used.

Phage vB_Eco_SLUR29 was found to rapidly lyse its host when used at an MOI of 1, but was less effective at lower MOIs. This was apparent in the local virulence index which is very close to zero at MOIs of 1×10^−6^ and 1×10^−7^ when the integrated area of the curve of infected and control samples are compared. To overcome this, we could have chosen a later point where lysis had occurred for cells infected at low MOIs. However, this would be well into the stationary phase of growth in the uninfected control so we chose to integrate from time zero to the onset of stationary phase as described in the original method ^19^. When compared to the virulence index of phages T7, T5 and T4, it has a lower virulence index of 0.37 ^19^ than any of these phages under any condition, suggesting it is not a particularly virulent phage.

A further observation of the properties of vB_Eco_SLUR29 infection, was the rapid recovery of infected cells. When infected at high MOIs there was rapid lysis of host cells, with recovery of cell numbers close to an un-infected control after 13 hrs. Whilst we did not explicitly test cells at the end of the killing assay, it is most probable that this recovery in cell growth is due to the rapid selection of resistant cells. The rapid development of resistance in *E. coli* to the closely related phage vB_Eco_S59 has been reported, suggesting there might be a minimal cost to developing resistance for this type of phage ^21^. However, the mechanism behind the development of resistance still remains to be elucidated.

The sequencing of vB_Eco_SLUR29 has expanded and helped to further clarify the phylogeny of phages within the genus *Tunavirinae*, with vB_Eco_SLUR29 a member of putative new genus that is clearly separate from the phage pSf-1 (Fig 4). Furthermore, the characterisation of phenotypic properties reveals that phages which are similar at the genomic level, have very different phenotypic properties. This highlights the need to assess the lytic properties of phage that are genetically similar, as it cannot always be assumed they will have similar infection properties. With increasing interest in the use of phages as therapeutics, the lytic properties are important factors that will need to be considered. In part, differences in lytic properties may result from the method used for carrying out one-step experiments. Therefore, we utilised the recently developed virulence index ^19^ which should allow more consistent comparison of the lytic properties of phages from different laboratories in the future.

## Acknowledgements

Bioinformatics analysis was carried out using MRC CLIMB Infrastructure MR/L015080/1. AM, JH, and SH were funded by NERC AMR -EVAL FARMS (NE/N019881/1). T.R. and S.M. were in receipt of PhD studentships funded by the Natural Environment Research Council (NERC) CENTA DTP

